# Using *fanyi* to reduce language barriers, promote information absorption, and communication

**DOI:** 10.1101/2023.12.21.572729

**Authors:** Difei Wang, Guannan Chen, Xiao Luo, Li Zhan, Shuangbin Xu, Qianwen Wang, Junrui Li, Rui Wang, Shaodi Wen, Guangchuang Yu

**Affiliations:** Nanfang Hospital, Southern Medical University, Guangzhou, 510515, China; Department of Bioinformatics, School of Basic Medical Sciences, Southern Medical University, Guangzhou, 510515, China; Wu Lien-The Institute & Department of Microbiology, Harbin Medical University, Harbin, 150081, China; Microbiome Medicine Center, Department of Laboratory Medicine, Zhujiang Hospital, Southern Medical University, Guangzhou 510515, China; Department of Oncology, The Affiliated Cancer Hospital of Nanjing Medical University & Jiangsu Cancer Hospital, Nanjing, 210009, China

## Abstract

Understanding the latest research advancements and sharing one’s own achievements is extremely important for young researchers. However, language barriers restrict the absorption of knowledge and career development. To address these issues, we have developed the fanyi package, which utilizes AI-driven online translation services to accurately translate between multiple languages. This helps researchers rapidly capture information, promotes the preservation of a multilingual environment, and ultimately fosters innovation in the academic community. Interpreting genes of interest (such as mutant genes) contributes to proposing molecular mechanisms, but accessing such information typically requires tedious manual retrieval. The fanyi package also provides tools for retrieving gene information from NCBI, enhancing the accessibility of gene information. Being able to quickly and efficiently obtain these pieces of information, coupled with translation capabilities, can help researchers better understand the information. The fanyi package is freely available via https://github.com/YuLab-SMU/fanyi. The translation functions are not limited to biomedicine research, they can be applied to any other domain as well.

## Introduction

English, as the first language of scientific research, plays an important role in disseminating scientific research results. However, this also poses challenges for non-native English-speaking researchers in writing and talking about their research. Many graduate students from non-English speaking countries face more or less obstacles to using English in the academic environment. Early-career researchers may find it challenging to master scientific writing in English in order to effectively convey the significance of their work, which can pose an extra obstacle when aiming to publish their work and advance their careers. In scientific writing, researchers often rely on various tools or language editing services. The application of large language models such as ChatGPT in academic writing has sparked considerable debate and remains controversial ^1^. This language barrier is also the reason why many local media and social platforms use their local language to interpret scientific results. With the development of artificial intelligence, text translation has become quite accurate. A large number of students use translation software to convert English materials into their language when reading scientific research papers and also use translation software to translate their domestic language into English when writing papers.

In biomedical research, a common scenario is for researchers to search for one term or gene after another, obtain explanations for the term or gene, and then use translation tools to translate this information into the local language, to quickly capture relevant information for interpretation. This is usually cumbersome and requires a lot of time. For example, when conducting single-cell analysis, the interpretation of marker genes in different cell clusters often requires the aforementioned method. To interpret these genes, as mentioned above, repeating operations for each gene requires a significant amount of time investment. Many researchers choose not to do this and instead directly use the genes for functional enrichment analysis ^2^. Only the results of enrichment analysis were used to selectively search for certain genes. Although it saves time, it is also very likely to miss some important genes.

In response to this situation, we have developed the R package, fanyi, which means “translation” in Chinese Pinyin. It can utilize online AI translation services to translate between multiple languages. At present, this package can support users to use Baidu, Bing, Youdao, Volcengine, Caiyun, Tencent, and ChatGLM translation services. With this package, users can translate the text using R language, which is widely used in multiple fields. For example, R is a necessary Swiss army knife for omics data analysis and visualization. With the help of fanyi, analysis results can be directly translated into the local language, and descriptions of functions, pathways, phenotypes, and diseases will become easier to understand. The fanyi package is freely available at https://cran.r-project.org/package=fanyi and https://github.com/YuLab-SMU/fanyi.

Retrieving gene information can be considered a daily task, such as searching for marker genes to better interpret cell types. To better serve researchers in the biomedical field with this translation tool, we have implemented automatic retrieval of gene descriptions. This allows for batch retrieval of analysis results, such as a list of differentially expressed genes, genes with high mutations in tumors, and marker genes for different cell types, etc. By automatically retrieving descriptions and summaries of genes from previous studies, it will greatly save the user time. Moreover, through translation, language barriers can be eliminated. Enable researchers to spend more time thinking about scientific questions, while fostering a multilinguistic culture in the science community.

## Results

### Fanyi retrieves marker gene information to assist in interpreting results

One of the main goals of biological research is to identify molecular mechanisms. After obtaining genes of interest, a significant amount of time is typically spent on searching and summarizing the functions of these genes to provide clues for proposing mechanistic hypotheses. The fanyi package can batch retrieve summary descriptions of genes from the NCBI database, greatly simplifying this step. Here, we use a single dataset from the Seurat example ^3^, containing 2,700 Peripheral Blood Mononuclear (PBMC) cells. After identifying the marker genes of all the cell clusters, we converted the gene symbols to ENTREZ gene IDs using clusterProfiler ^2^ and then queried gene information using the gene_summary function provided in the fanyi package.

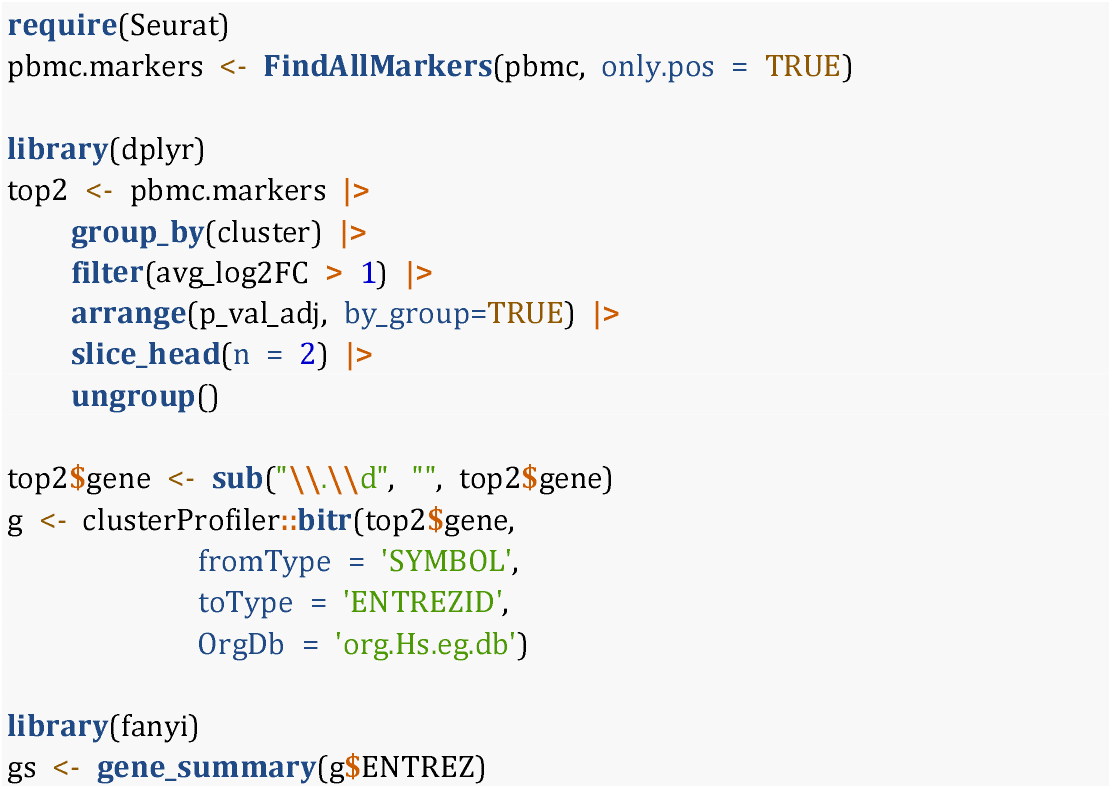

As demonstrated in Figure 1, the information output by FindAllMarkers can be integrated with the output of gene_summary, and all these pieces of information can be exported as a table to help the user understand the marker genes. Combining expression levels, p values, and detailed gene function descriptions of marker genes, researchers can better understand and interpret the cell types and functions of different cell clusters.

**Figure 1.**
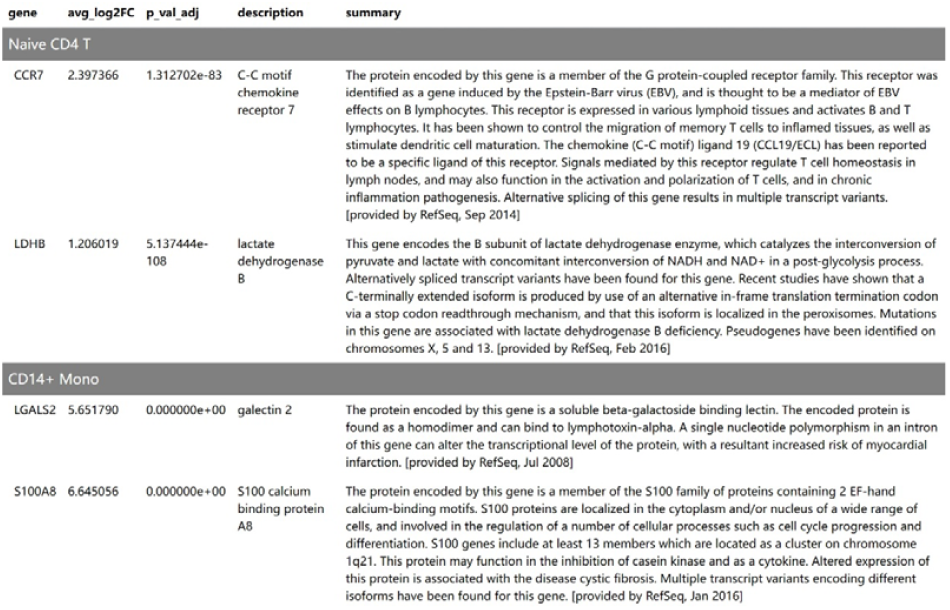
Table of marker gene information. As a demonstration, only the top two marker genes belonging to two cell clusters (Naïve CD4 T and CD14+ Mono) are displayed.

### By combining information/knowledge, fanyi helps create easy-to-understand infographics

Infographics contain data visualization such as bar charts and descriptive text representing information, making the conveyed information clearer and more quickly understood. The gene description and summary information queried by the gene_summary function can be used in data visualization to provide a more informative graph.

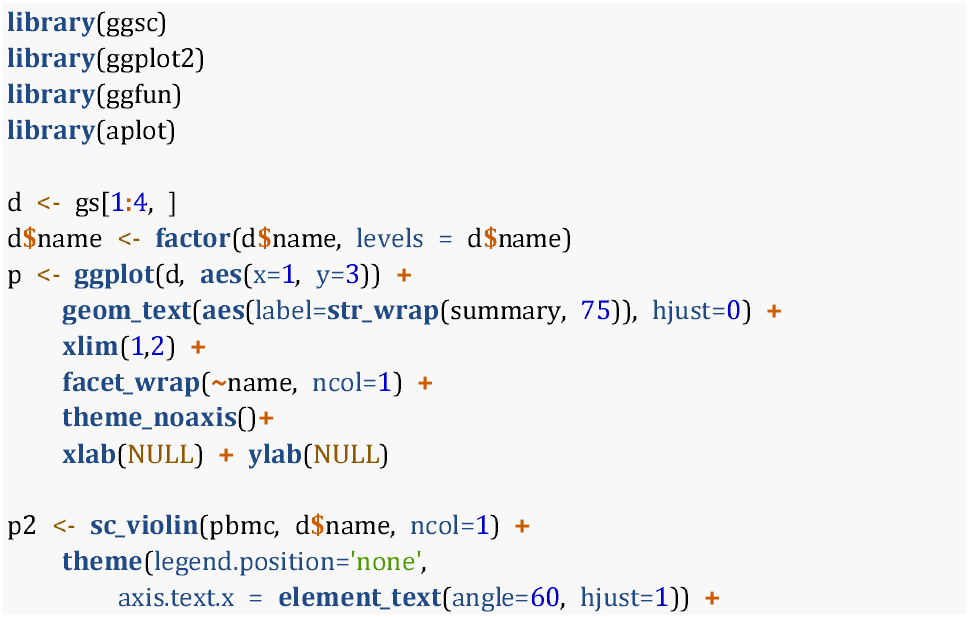
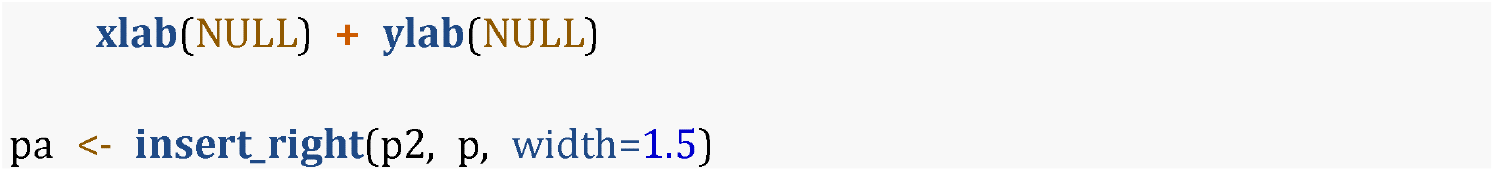

The above source codes employ the in-house developed package, ggsc, to visualize the expression distribution of the selected marker genes among different cell clusters as violin plots and display the gene summary side by side to help the user interpret the results as demonstrated in Figure 2A.

**Figure 2.**
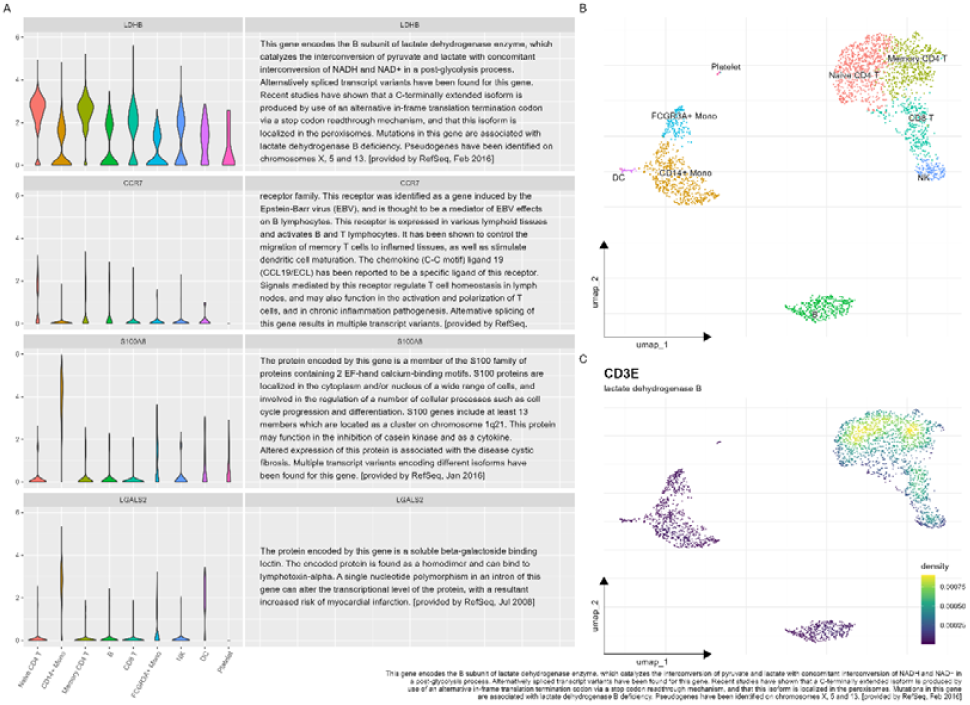
Use gene information to create an informative graph that contains a visualization of data and biological knowledge. (A) Violin graphs display gene expression distributions among different cell clusters with gene summary information side by side to explain the gene functions. (B) Cell type annotation on a UMAP plot. (C) Density plot to visualize the expression levels of the CD3E gene across different cells on a UMAP with the information of full gene name and summary of gene function.

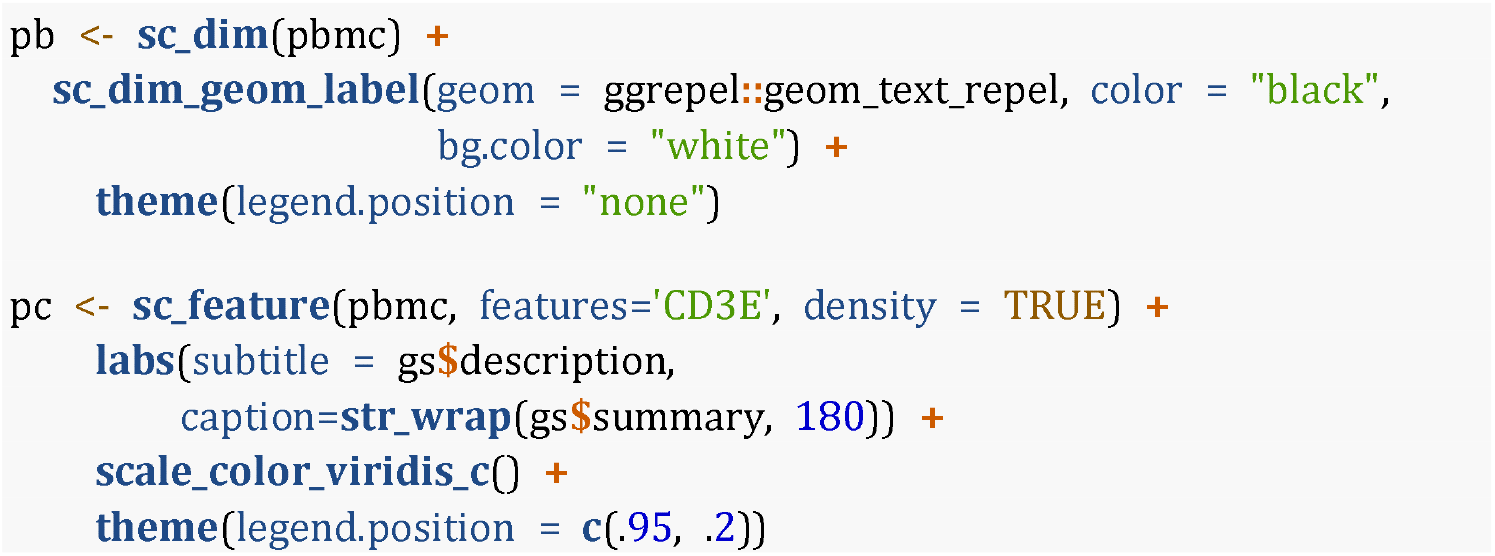

We further use ggsc to visualize the pbmc dataset. Figure 2B illustrates the cell clustering results and Figure 2C visualizes the expression of the CD3E gene in different cells as a density plot. With the gene information provided, we can display the gene description as a plot subtitle and the gene summary information as a plot caption. The information provided in Figure 2C helps us to capture the idea of how the CD3E gene functions and to interpret the expression distribution among different cell clusters.

### Fanyi uses AI-driven translation services to help researchers quickly digest information

The language barrier makes it difficult for professional content to be quickly understood and absorbed, while also hindering researchers from effectively expressing and conveying information. The fanyi package, with the assistance of multiple AI-driven translations, can effectively perform batch and automated conversion of information across multiple languages. In this example, we used an RNA-Seq dataset of four human airway smooth muscle cells (GEO: GSE52778) ^4^. This dataset can be accessed via the Bioconductor package, airway.

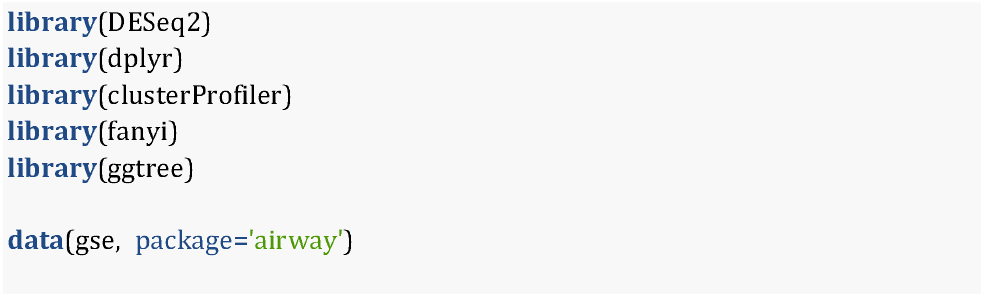
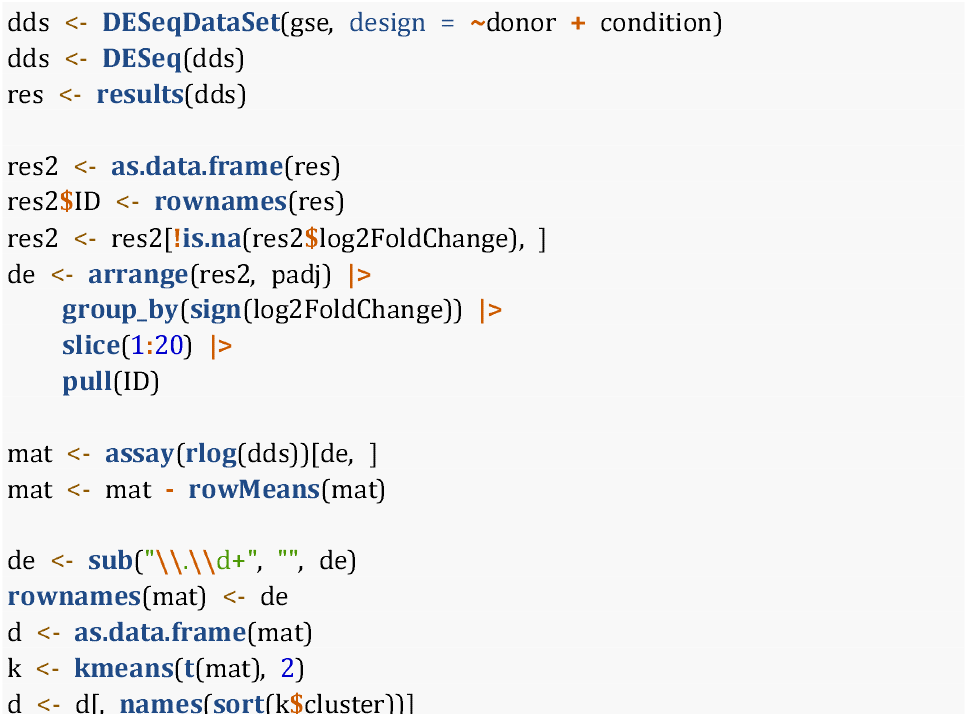

As shown in the above code, we employed DESeq2 ^5^ to identify differentially expressed genes. We extracted 20 genes from up-regulated and down-regulated genes respectively. These 40 genes were further used to demonstrate how fanyi can help interpret the results.

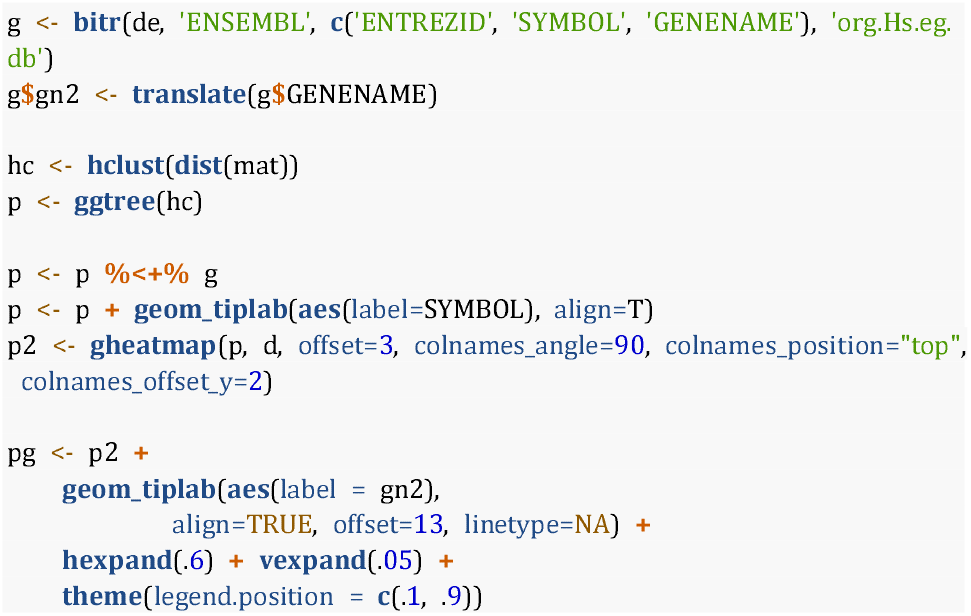

We used bitr function to convert gene IDs and used the translate function to convert the full gene name into Chinese. We then applied the hierarchical clustering method to cluster the genes based on the expression profiles. The clustering tree was visualized using ggtree ^6–8^ with gene symbols and then gheatmap was utilized to visualize the tree with the expression matrix ^9^. Finally, we displayed the Chinese version of the full gene names to help users better understand the functions of these differential genes (Figure 3A). With fanyi, users can translate information into their local language with more than two hundred languages supported and display the information with the English version side by side. We no longer need to translate professional vocabulary manually. It saves a lot of time and helps users to quickly capture the information when looking at the figure.

**Figure 3.**
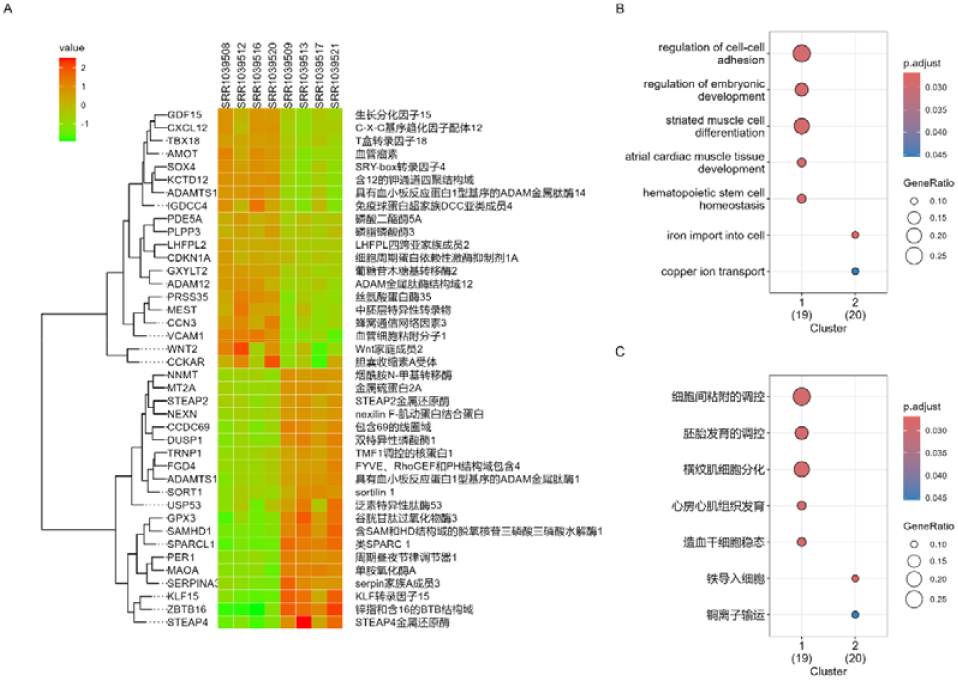
Translated information in the local language helps users quickly capture and understand the information presented. (A) A heatmap visualizes differential expressed genes across different samples with gene symbols and translated full gene names. (B) Functional characterization of differentially expressed genes with different expression patterns. (C) A translated version of (B).

As demonstrated in Figure 3A, the genes can be divided into two clades, we applied cuttree function to separate the genes into two clusters and then employed clusterProfiler ^2,10^ to characterize biological processes regulated by these two gene clusters as demonstrated in Figure 3B. In fanyi, we provide a helper function, translate_ggplot, to help users translate words and sentences in a ggplot graph. As demonstrated in Figure 3C, the biological process terms presented in Figure 3B are translated into Chinese with only one line of function call.

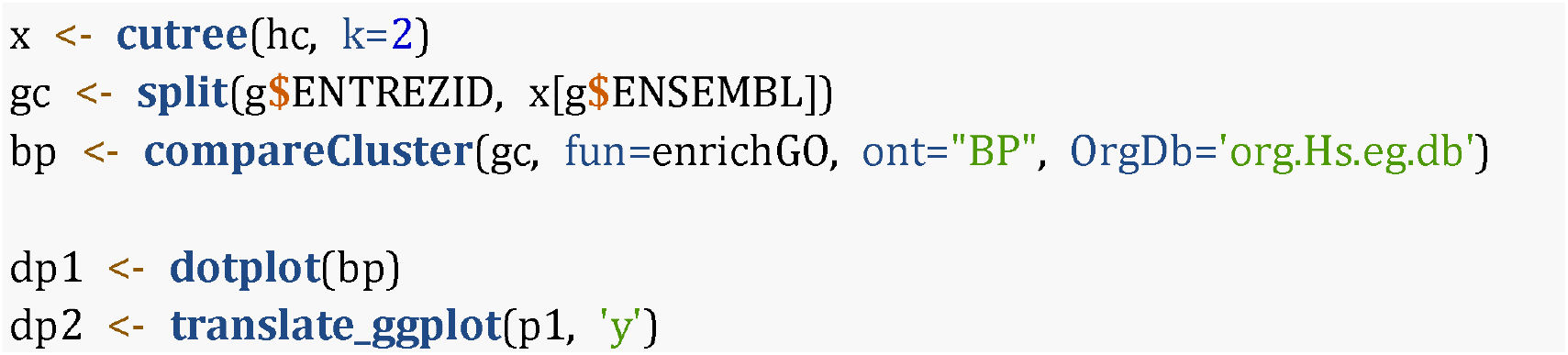

## Discussion

Language barrier has serious consequences for science and science education, including inequality for under-represented communities, difficulties in disseminating non-English knowledge, and challenges in uptaking knowledge in educational settings ^11^. Language shapes the way we think ^12^. People speaking different languages tend to think in different ways, which can bring new perspectives and ideas. The diversity of languages helps foster innovation and creativity, which is exactly what the academic community needs.

The fanyi package helps lower the language barrier by employing various AI-driven online translation services. In particular, information can be translated into the local language to help quickly uptake the information, non-English-language information can be translated into English to increase the visibility of the content, and existing resources and new discoveries can be disseminated in multiple languages to communicate effectively to non-English speakers. This will promote the preservation of a multilingual environment in the academic community.

The fanyi package presented here can also help retrieve and interpret biological information. It not only makes the gene information accessible but also saves researchers a significant amount of time.

Querying gene information can be used to quickly create a summarized table with rich information for marker genes, differential expression genes, highly mutated genes, etc., to help users digest how the genes work. The information query by the package can also be utilized to annotate the plot to create an informative graph with more or less self-explanation.

## Acknowledgments

This study was supported by the National Natural Science Foundation of China [32270677]. The manuscript was drafted with the help of the fanyi package. We appreciate the feedback and support from fanyi users.

